# MetaConnect, a new platform for population viability modelling to assist decision makers in conservation and urban planning

**DOI:** 10.1101/2019.12.30.890921

**Authors:** Sylvain Moulherat, Elvire Bestion, Michel Baguette, Matthieu Moulherat, Stephen C.F. Palmer, Justin M.J. Travis, Jean Clobert

## Abstract

In a context of global change, scientists and policy-makers require tools to address the issue of biodiversity loss. Population viability analysis (PVA) has been the main tool to deal with this problem. However, the tools developed during the 90s poorly integrate recent scientific advances in landscape genetics and dispersal. We developed a flexible and modular modelling platform for PVA that addresses many of the limitations of existing software. MetaConnect is an individual-based, process-based and PVA-oriented modelling platform which could be used as a research or a decision-making tool. Here, we present the core base modelling of MetaConnect. We demonstrate its potential use through a case study illustrating the platform’s capability for performing integrated PVA including extinction probability estimation, genetic differentiation and landscape connectivity analysis. We used MetaConnect to assess the impact of infrastructure works on the natterjack toad metapopulation functioning.

## Introduction

In a context of rapid global change, habitat loss and habitat fragmentation have become the major threats to biodiversity (IUCN 2013), and have been considered for some time as the principal cause of species extinction (Dobson, Bradshaw & Baker 1997; Millenium Ecosystem Assessment 2005). They have consequences from the ecosystem (Fahrig 2003; Cardinale *et al.* 2012; de Mazancourt *et al.* 2013) to the genetic scale (Ingvarsson 2001; Baguette *et al.* 2013) by modifying landscape patterns in a four-step process: reduction in amount of habitat, increase in number of habitat patches, decrease patch sizes and increase in patch isolation (Fahrig 2003). This alteration of landscape patterns has diverse effects on population dynamics (e.g. Baguette *et al.* 2013). As patches become smaller, the size of the population supported decreases; this can increase stochastic risk of extinction from demographic processes (Legendre *et al.* 1999; Reed *et al.* 2002), but also from genetic stochasticity: small populations are more subjected to risk of inbreeding and consanguinity depression (Brook *et al.* 2002b), loss of genetic diversity and mutational accumulation (Rowe & Beebee 2003), leading to the extinction vortex (Gilpin & Soulé 1986; Fagan & Holmes 2006). Moreover, by increasing inter-patch distances and therefore dispersal cost, habitat loss and fragmentation decrease the probability of individuals moving between sub-populations. This can hinder recolonization or the demographic rescue of patches where sub-populations have become extinct, or declining, potentially leading to stochastic extinction of metapopulations (Fahrig 2003). Furthermore, by reducing gene flow between sub-populations, isolation can lead to genetic differentiation of sub-populations and impede genetic rescue of highly inbred sub-populations (Ingvarsson 2001; Keller & Waller 2002; Tallmon, Luikart & Waples 2004).

Historically, an ecological tool much used by scientists and conservation managers was population viability analysis (PVA), which aims at estimating extinction or quasi-extinction probability of a species (Boyce 1992; Legendre & Clobert 1995; Beissinger & Westphal 1998) and which was used to inform conservation programs (Southgate & Possingham 1995; Ferriere *et al.* 1996; Letcher *et al.* 1998; Schtickzelle & Baguette 2004; Radchuk *et al.* 2013). Most of the PVA models focused on species population dynamics (Lindenmayer *et al.* 1995; Brook *et al.* 1999; Legendre *et al.* 1999), and offered limited flexibility regarding population genetics (but see Lacy, Borbat & Pollak 2009) or metapopulation functioning (*Reed et al.* 2002; Pe’er *et al.* 2013). PVA is principally focused on demographic and genetic processes and individual movement behaviour determining species persistence (Keller & Waller 2002; Fahrig 2003; Piou & Prevost 2012; Frank & Baret 2013; Noel, Machon & Robert 2013). Population genetics has made major advances during recent decades, and it is now possible to identify at generation *g* those individuals that are offspring of individuals immigrating at generation *g-1*; and assign these immigrants to their original population (Beerli & Felsenstein 2001). Besides, the rise of landscape genetics allows the assessment and quantification of how landscape elements affect gene flow in a metapopulation (Manel *et al.* 2003; Manel & Holderegger 2013). Integrating both population and landscape genetics approaches in models of metapopulation functioning to support conservation managers’ and policy makers’ plans should be highly valuable (Baguette *et al.* 2013; Pe’er *et al.* 2013).

The growth of a community of ecological modellers using individual-based models (IBMs) rather than the mathematical approaches has led to the production of a huge number of models (DeAngelis & Mooij 2005) and metrics (Moilanen & Nieminen 2002; Calabrese & Fagan 2004). Moreover, most such models were developed to answer specific questions, which renders comparison between outputs difficult, if not impossible (DeAngelis & Mooij 2005; Kindlmann & Burel 2008; Pe’er *et al.* 2013), and their application inefficient out of the narrow context for which each was typically developed (Grimm *et al.* 2004; Kindlmann & Burel 2008). Generic PVA modelling platforms built from IBM and process-based modelling are lacking (but see Grimm *et al.* 2004; Lacy, Borbat & Pollak 2009; Bocedi *et al.* 2014) or do not permit a sufficient level of flexibility (VORTEX is spatially implicit, RangeShifter and MetaX have limited demographic modules) to deal with a large spectrum of conceptual framework and ecological themes (DeAngelis & Mooij 2005; Evans, Norris & Benton 2012; Evans *et al.* 2013; Purves *et al.* 2013).

Because PVA is a relevant basic decision-making tool (Brook *et al.* 2000; Brook *et al.* 2002a; Pe’er *et al.* 2013), we developed MetaConnect, a generic PVA-based IBM following the Beissinger & Westphal (1998) framework, which allows the design of a range of models from very simple to highly detailed and which integrates demography and genetics in a spatially-explicit context (Baguette *et al.* 2013). MetaConnect not only aims at performing traditional PVAs based on demographic data, but also integrates the recent development of population and landscape genetics, which allows the assessment of functional connectivity (sensu Taylor, Fahrig & With 2006). We expect that this integrated modelling platform will be a useful tool for scientists, conservation managers and policy makers. Here we present MetaConnect’s core base modelling, its validation (Appendix A) and present a case study demonstrating its use to assess the impact of infrastructure development on the population viability, genetic differentiation and functional connectivity of existing populations of the endangered natterjack toad (*Epidalea calamita*).

## Model

### Model design

MetaConnect simulates metapopulation dynamics and genetics using the species life cycle and life history traits, the landscape characteristics and their interactions. The simulations allow inferring of individual dispersal and local and global extinction probabilities, genetic diversity and genetic differentiation (from classical Fst analyses or as input files for the Structure software (Pritchard, Stephens & Donnelly 2000)).

MetaConnect is an individual- and process-based model which means that:

- all individuals in the model are independent and behave in respect to their phenotype.
- patterns emerging in the different outputs of the model are the products of flexible and adjustable rules implemented in the model.

#### Model structure

##### Landscape

The landscape is imported as two shapefiles and then rasterized by MetaConnect.

- Patches: locates suitable habitats for the focal species. The carrying capacity (inidividual/m²) can be assumed to be constant, or an optional shapefile of carrying capacities can be provided.
- Costs: provides a coefficient (of rugosity) representing the ability of a given species to move through each habitat type of the landscape. The higher the cost, the harder to cross.

##### Demography

Population dynamics is represented by a succession of individual states linked by transitions. The user builds the species life-cycle by assembling “bubbles” representing the individual state and “arrows” representing transition rules between individual states (Figure 1, 2). The “bubbles”, hereafter regarded as classes, correspond to age classes, sex or anything that can be defined as a group of individuals with the same demographic characteristics. Density dependence can be scramble or contest and designed as a part of transitions. The mating system can be chosen from monogamy, polygamy, polyandry and/or polygyny (*Legendre et al. 1999*). The demographic parameters (Table 1) can be patch-specific. Environmental stochasticity has been included as random processes inducing normal variation around the patch’s mean value of demographic parameters truncated to realistic values set by the user. As an example, the fecundity parameter follows a Poisson distribution (demographic stochasticity) with parameter λ equal to the average fecundity 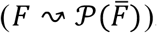. The average fecundity can vary from one patch to another and within simulation time steps following a Gaussian distribution (Table 1).

**Figure 1:**
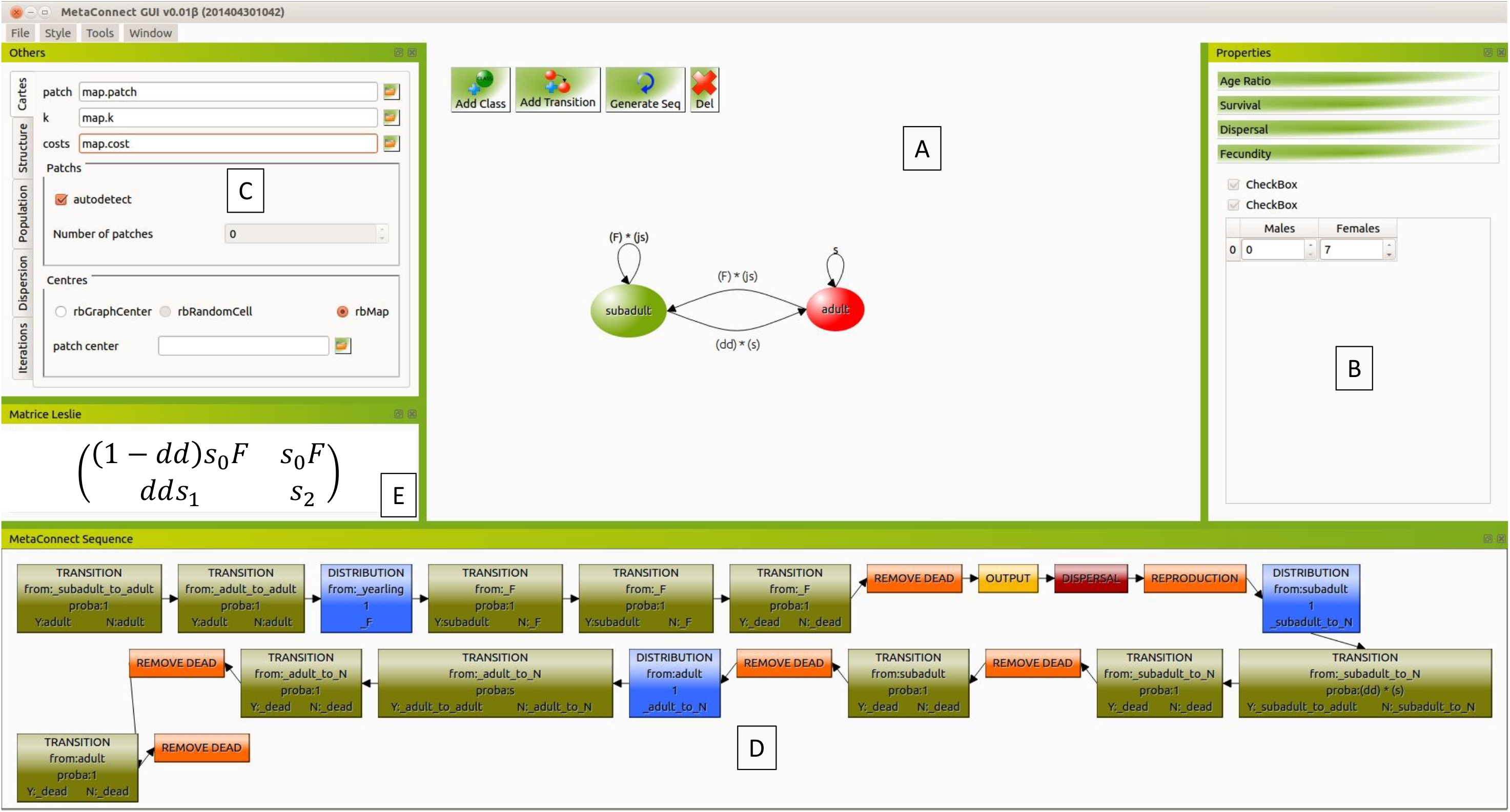
Screenshot example of the Legendre et al. (1999) passerine life cycle with two age-class and two sexes that can be modelled with MetaConnect. The user defines the species life-cycle as a combination of “bubbles” (add class) and “arrows” (add transition) (A). The species life history traits are set up in the B section and the run setting is defined in the C. Then, the MetaConnect workflow (D) and the Leslie matrix (E) are generated automatically.

**Figure 2:**
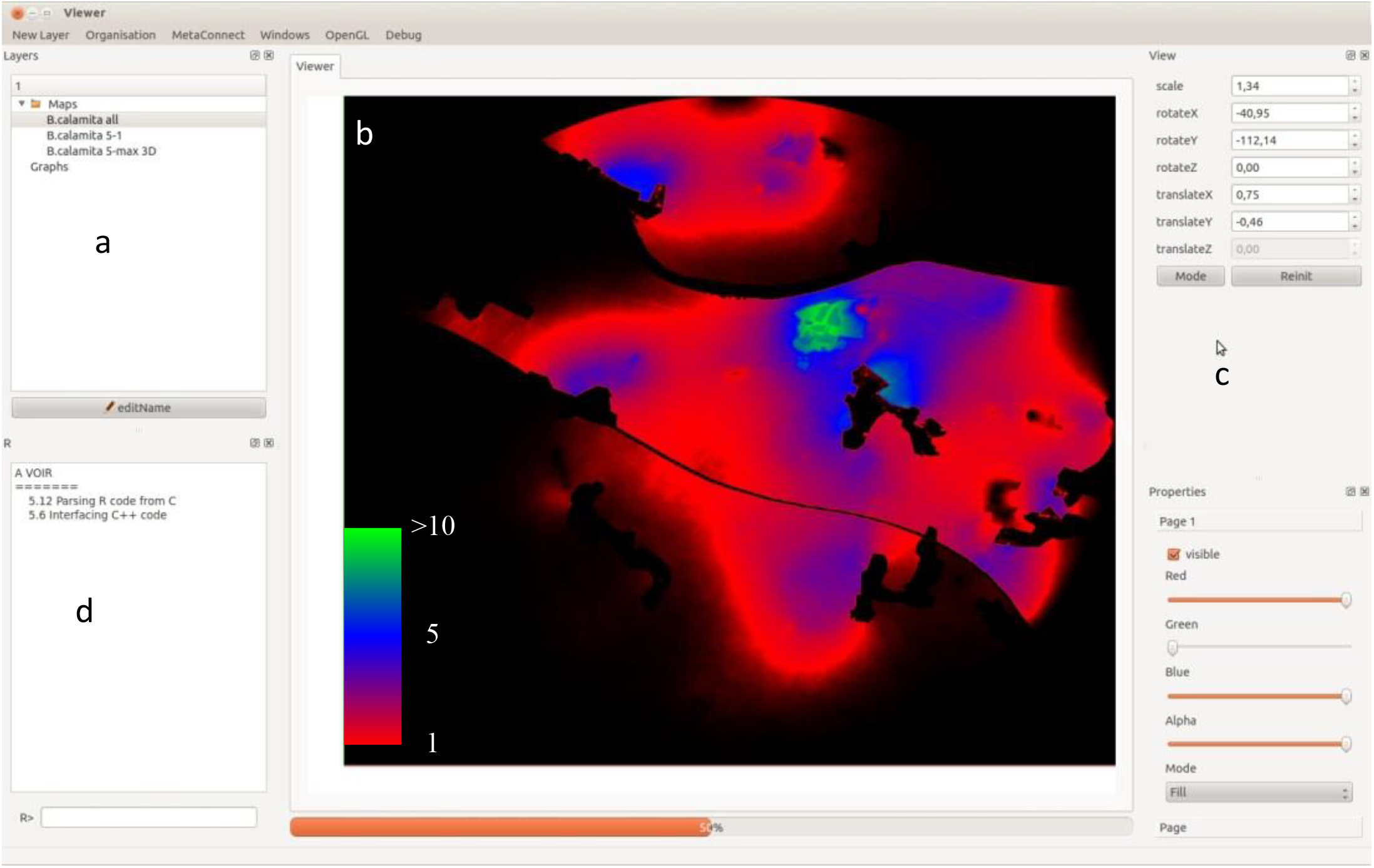
*E. calamita* landscape use during dispersal (indiv./year) estimated with MetaConnect. Data to be displayed or analysed are selected in section *a* and displayed in *b* in two or three dimensions. Section *c* allows setting of the display options, and *d* is an R console to perform simple analysis.

**Table 1:** Nomenclature of MetaConnect’s main parameters and variables.

| Parameters and variables | Description | Distribution law of random variables |
| --- | --- | --- |
| *Demographic characteristics* |  |  |
| $K$ | Carrying capacity* | |
| $k$ | Total competition coefficient | |
| $s_i$ | Survival of individual from class $i$ | Bernoulli |
| $F$ | Fecundity | Poisson |
| $\sigma$ | Primary sex ratio | Bernoulli |
| *Mating system* |  |  |
| Mating system | Mating system assumptions |  |
| $H_m$ | Male harem size | Poisson |
| $H_f$ | Female harem size | Poisson |
| *Genetics* |  |  |
| $L$ | Number of loci | |
| $A$ | Number of alleles per locus | |
| $\mu$ | Mutation rate | |
| *Dispersal* |  |  |
| $p_d$ | Dispersal probability | Bernoulli |
| Dispersal rule | Dispersal algorithm |  |
| *Initialization and model parameterization* |  |  |
| $N_0$ | Initial number of individuals | |
| $\sigma_0$ | Initial sex-ratio | |
| $f_a$ | Initial class structure | |
| $T$ | Time steps | |
| $MC_l$ | Number of landscape random generations | |
| $MC_d$ | Number of population dynamic simulations per landscape | |

\* The carrying capacity $K$ is derived from the competition coefficient and the competition assumption.

Dispersal probability *p*_*d*_ is implemented by setting a proportion of individuals in a given class leaving a patch. The density-dependent recruitment probability *p*_*r*_ is determined by equation 1 where *N*_*T*_ can be a chosen combination of the number of individuals per class (i.e. *N*_*T*_.could be the total population or the number of individual in a given class) (Caswell 2001).

Equation 1:

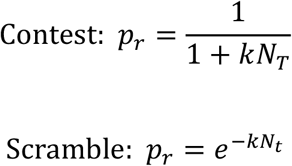

Dispersal is age- and sex-dependent, and the process by which individuals disperse can be chosen from three families of movement rules:

- Dispersal between patches is modelled by a probability for an individual to reach another patch, ignoring rugosity coefficient. The probability of reaching a patch can be equal between patches, depend on the Euclidean distance between patches’ centres, or be set manually.
- The interaction between individuals and their environment depends on rugosity coefficients. This family comprises a random-walk (RW) and a correlated random-walk rule (CRW). The CRW assumes a degree of directional persistence, (i.e. movement direction at time *t+1* depends on the direction taken at time *t*) and not solely an environmental based one. Dispersing individuals benefit from a “energy gauge” decreasing at each step of the (C)RW in respect of the cell cost weighted by direction (1 for straight moves, 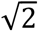 for diagonal moves).
- This family of rules assumes that individuals have knowledge of their environment, and move by one of two methods. From a focal patch, the least cost path (LCP) algorithm usually assumes that a single patch can be reached (Botea, Müller & Schaeffer; Adriaensen *et al.* 2003; Pe’er & Kramer-Schadt 2008; Barraquand, Inchausti & Bretagnolle 2009). Such an assumption is unrealistic, and to relax it we implemented a multiple LCP movement rule, in which we calculated all possible LCPs between the focal patch and all other patches (Urban *et al.* 2009; Foltete, Clauzel & Vuidel 2012). Then, for reachable patches (i.e. LCP length less than the maximum dispersal distance), the probability to reach a patch is inversely weighted by the LCP length (number of map cells crossed) or cumulative cost (total cost of all map cells crossed). We also adapted the Stochastic Movement Simulator (SMS) (Palmer, Coulon & Travis 2011; *Aben et al. 2014*; Palmer, Coulon & Travis 2014; Coulon *et al.* Submitted), which relaxes the assumption of omniscience inherent in the LCP approach. With the SMS rule, individuals make movement decisions based on the environment within a limited perceptual range and a tendency to directional persistence similar to that in a CRW. At each movement, the SMS algorithm calculates a movement probability for each cell surrounding the current cell based on the rugosity coefficient of the cells in the perceptual range (see Palmer, Coulon & Travis 2011 for details).

The dispersal event ends when the individual dies or reaches a patch different from his original patch regardless of the arrival patch quality.

##### Genetics

Individuals are genetically tagged using neutral polymorphic loci. The number of loci and number of alleles per locus can be specified by the user. A single mutation rate (a probability of creating a new allele without possibility of reverse mutation) implemented in the model allows the production of new alleles at each locus during simulations, and can be specified by the user. Gene transmission is assumed to be Mendelian and siblings are assumed to have the same father (randomly chosen from the female harem for polyandrous cases).

#### Model outputs

The model provides many forms of outputs based on focal species life history traits and landscape maps, which are adaptable to various theoretical and applied contexts. The outputs report the results at three levels at a frequency specified by the user, allowing dynamic visualization of the simulations:

- Demography: population size is split into the classes and sexes implemented in the model. The model derives extinction probability, colonization probability and time to extinction and colonization. These indicators are calculated at the local (patch) and global (metapopulation) scales.
- Dispersal: the model provides the number of individuals reaching a new patch or dying during dispersal. Maps of cell occupancy are drawn from successful dispersal events (number of individuals visiting each cell during the whole run).
- Genetics: genetic diversity and differentiation (Fst, Fis, Fit, He and Ho) at the local and/or global scale applicable to each statistic.

All these outputs can be directly plotted using the MetaConnect project manager (figure 2) or extracted as text files. In addition R (R Development Core Team 2005) has been incorporated to the project manager, which allows direct analysis of the outputs.

### Model sensitivity

MetaConnect is a highly flexible and modular IBM, which means that dozens of variables can be specified in various modelling contexts, rendering global sensitivity analyses impossible to run (Cross & Beissinger 2001; Naujokaitis-Lewis *et al.* 2009; Pe’er *et al.* 2013). A thematic sensitivity analysis will be presented in the MetaConnect user manual, in which sensitivity of extinction probabilities, genetic structure and connectivity metrics will be analyzed in relation to the appropriate model parameters and their relative contribution to the sensitivity estimate (Cross & Beissinger 2001).

## Case study: Changes in the metapopulation functioning of an existing natterjack toad population under scenarios of economic development

### MetaConnect parameterization

We used a model designed with MetaConnect to determine the potential impact of the development of an industrial area and a terrestrial transport infrastructure (high-speed railway), both alone and together, on the population viability of *Epidalea calamita* populations in south-western France close to Agen (44°11’36”N, 0°31’14”E) (figure 3).

**Figure 3:**
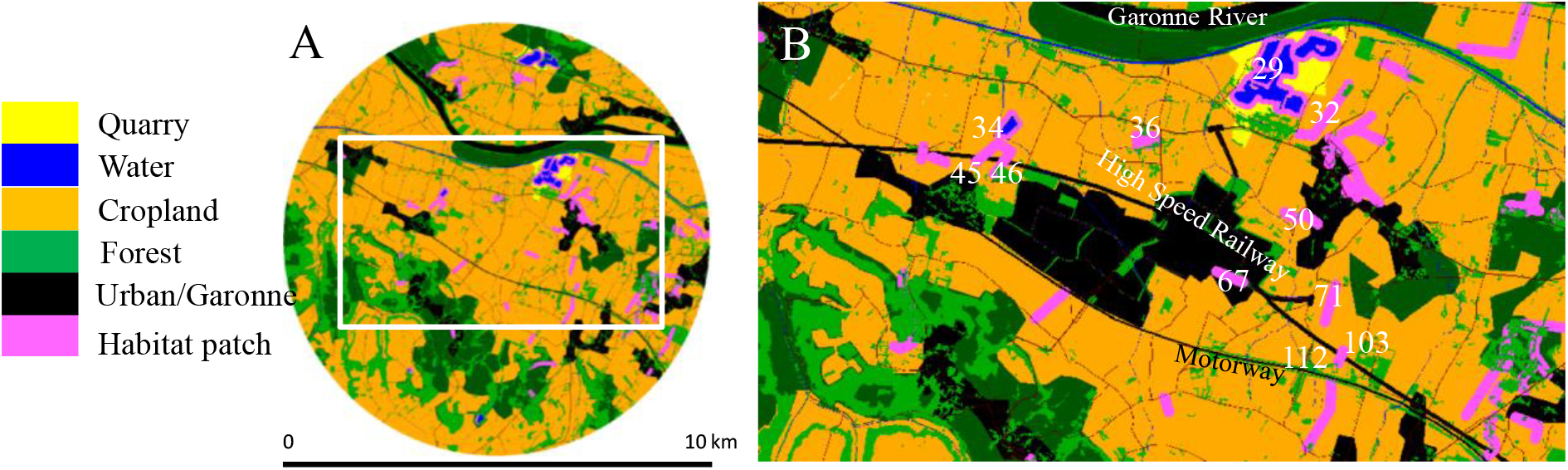
Site of industrial and transport development in south western France close to Agen. Panel A presents the complete study site used for MetaConnect simulations, rasterized to a grid of 10m × 10m cells. Panel B shows enlarged the future development of an industrial area and of a high-speed railway. Our analysis is focused on the patches numbered in white. Patch 67 will be destroyed by the industrial area development. Patches 45, 46 and 112 do not currently exist, but will appear with the building of the high-speed railway as a result of the fragmentation of respectively patches 34 (split into 34, 45 and 46) and 103 (split into 103 and 112).

Habitat patches were determined *a priori* based on expert assessment (figure 3). Preferences for landscape elements were derived from those experimentally determined by Stevens et al. (2006). *E. calamita* was assumed to disperse following the stochastic-movement-simulator (SMS) assumptions (Palmer, Coulon & Travis 2011; Coulon *et al.* Submitted). MetaConnect was parameterized with published values of life history traits (Stevens & Baguette 2008) (Appendix B).

We analysed MetaConnect genetic outputs for all the patches containing more than 10 individuals at the end of each run of 100 time steps. This analysis was performed using STRUCURE (Pritchard, Stephens & Donnelly 2000) with an admixture model assuming that the origin population of an individual is known and the allele frequencies are independent. STRUCTURE runs were performed for a variable number of clusters between 1 and 7, and with 10 iterations of 100000 steps (50000 burn-in and 50000 analysis steps to ensure model convergence). This procedure was reproduced for each iteration of the simulation of a given scenario. We determined the best number of genetic clusters following the Evanno method (see Evanno, Regnaut & Goudet 2005) using STRUCTURE HARVESTER (Earl & Vonholdt 2012). We counted the assignation of a patch to a cluster for each MetaConnect run analysed with STRUCTURE and tested the clustering robustness by performing a χ² test per patch.

### Current metapopulation functioning

Within the study site, (10 km around the industrial area development), the *E. calamita* population is not threatened (extinction probability *p*_*e*_ = 0) and we observed two main dispersal corridors. The major corridor joins the north (quarry, patch 29 and 32) to the south (motorway, patch 103) and is stopped by the motorway in its southern part and by the Garonne River at its northern part. The minor corridor joins the quarry (east) to a pond (patch 34, west) (figure 4.A). Our focal area covers these two corridors (figure 3). Both corridors incorporate stepping-stone sites (North-south: patches 50 and 67, east-west: patch 36) (figure 4.A). Within the focal area all the patches have a very low extinction probability (see figure 4), and except for patch 103, all patches are able to exchange individuals with all the others. Indeed, only patch 103 is unable to provide individuals to patches 34, 36, 29 and 32 or to receive individuals from patches 34 and 36.

**Figure 4:**
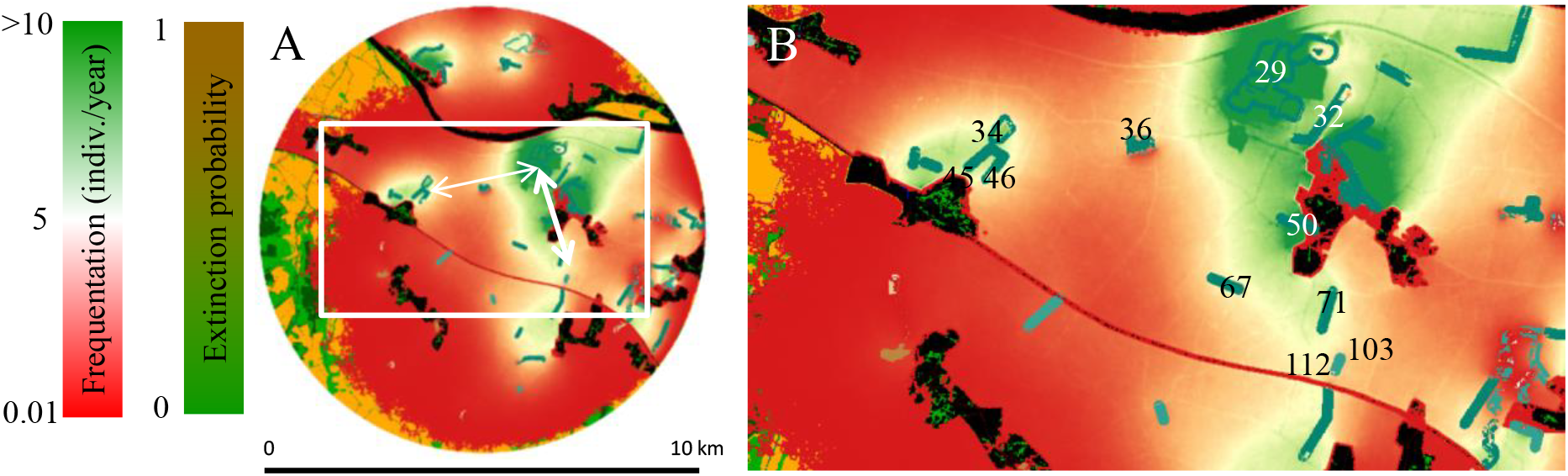
MetaConnect output layers of patch extinction probabilities and mean cell frequentation during efficient dispersal events. At the global scale, dispersing individuals follow two main corridors (white arrows in panel A): a major one along the north-south axis and a minor one along the east-west axis.

The analysis of the genetic output suggests that currently the study site is divided into three separate clusters (all cluster differentiation p.values<0.05). The first one is situated at the north of the Garonne, the second between the Garonne and the motorway and the third at the south of the motorway (figure 5.A).

**Figure 5:**
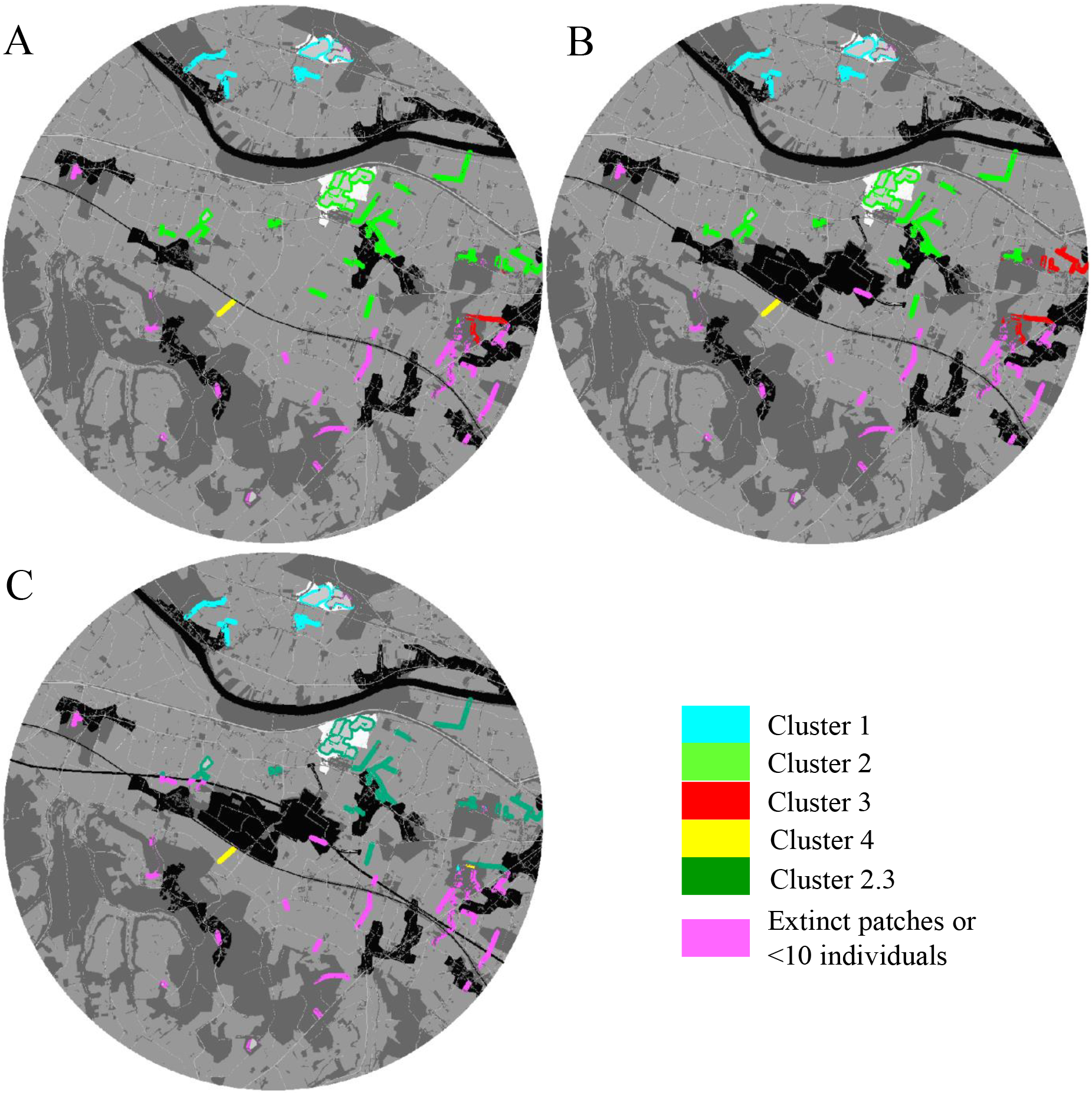
Genetic clustering of the suitable patches for *E. calamita*. The development of the industrial area (B) does not change the genetic structure of the *E. calamita* population from the current situation (A). 4 clusters are identified and separated by the Garonne (isolating clusters 1 and 2), a forest (barrier between cluster 2 and 3) and the motorway (separating cluster 2 from 4). The addition of the high-speed railway (C) would modify the genetic exchanges and only two clusters would be kept (cluster 1 and 4). The 2.3 cluster is neither aggregated to the cluster 1 nor 4 (all p.values > 0.05) due to the reduction of the number of individuals per patches and the change in the inter-patch genetic exchanges.

### Expected consequences of the industrial area and of the high-speed railway building

The industrial area development and the associated destruction of patch 67 leads to a reduction of individual flow between patches to the north and south of the industrial area and to a population size reduction within the southern patches (figure 6.B). Such a modification would not threaten *E. calamita* persistence in the study site (all *p*_*e*_ = 0, figure 6.D). However, dispersal would be more concentrated between patches 29, 32 and 50 due to the repelling effect of the industrial area for *E. calamita* (figure 6.D). In addition, the reduction of dispersal along the north-south axis should lead to a fourth genetic cluster in the western part of the study sites (Figure 5.B).

**Figure 6:**
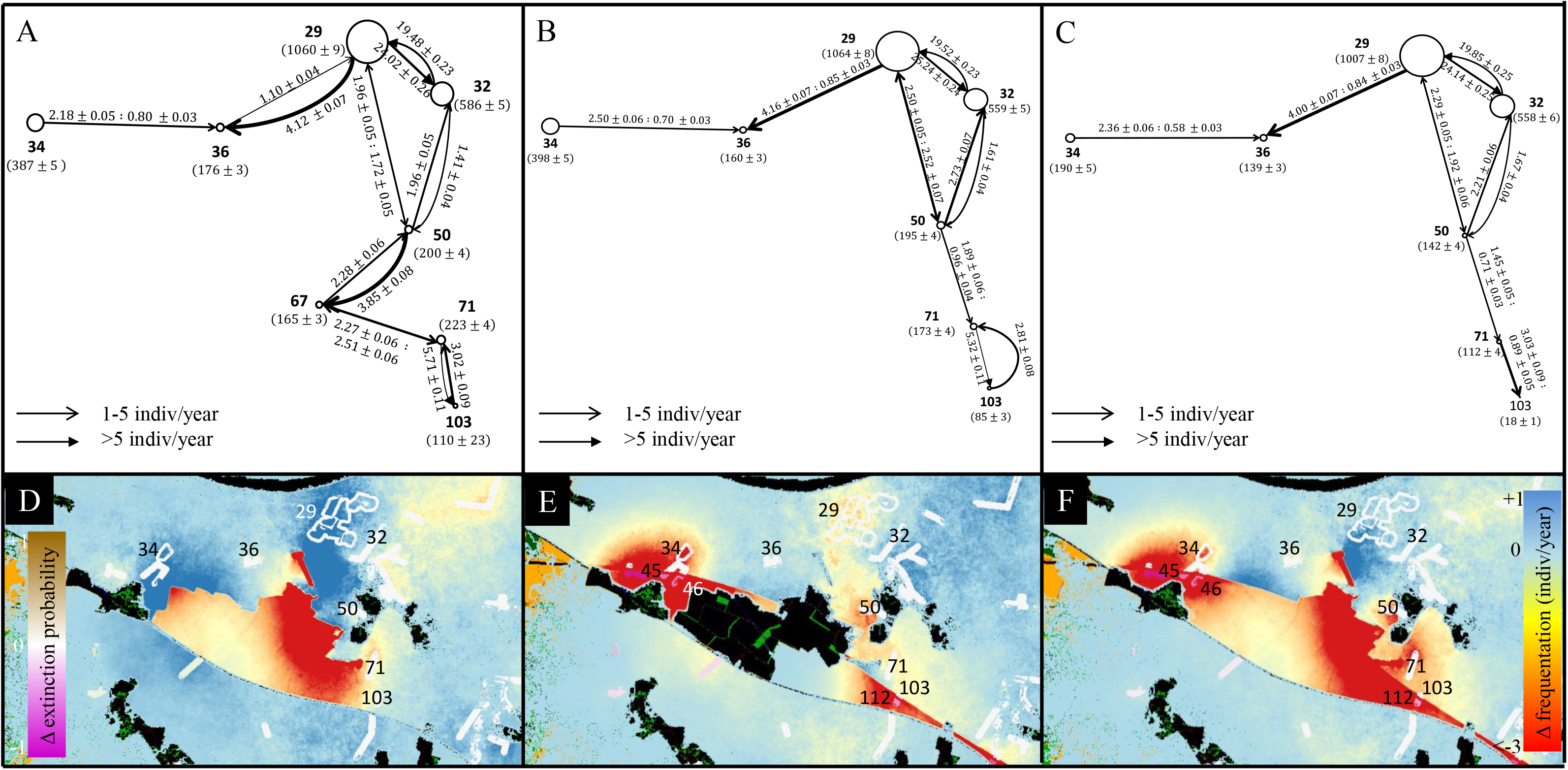
Graphical representation of the focal area metapopulation functioning derived from MetaConnect outputs for the current situation (A) with the industrial area (B) and with both the industrial area and the high-speed railway (C). Node sizes are proportional to patch population sizes (mean ± SE). Arrows represent dispersal intensity (mean ± SE) and direction (bidirectional arrows values correspond to smaller patch number to larger patch number flow: larger patch number to smaller). Maps represent the change in extinction probabilities per patch between scenario B and scenario A and similarly the change in cell occupancy during efficient dispersal. Panel D corresponds to the metapopulation functioning variations after the development of the industrial area. Panel E presents the variation between the situations after the industrial area development and after the high-speed railway building. Panel F summarizes the cumulative differences from the current metapopulation functioning to the expected functioning after the development of the industrial area and the high-speed railway.

The high-speed railway project will fragment the study site landscape. This would lead to the reduction of patches 34 and 103 areas (since none of the newly created patches 45, 46 nor 112 could shelter a population (Respectively *p*_*e*_ = 0.71, 0.71, 0.61, figure 6.E)) inducing the reduction of their sheltered population sizes (figure 6.C). In addition, dispersal along the north-south corridor is reduced to the south of the patch 50 (figure 6.C). Moreover, the connectivity along the north-south corridor is also decreased due to the population size reduction of patches 71 and 103. The reduction of the patch population size and the change in the connectivity along the north-south axis change the clustering outcome of these last simulations. Indeed, if the Evanno method suggests that 2 clusters can be identified, the χ² test shows that the assignation to a second cluster does not differ significantly from a random assignation, suggesting that only a single cluster exists or that the cluster 1 (north of the Garonne) is poorly differentiated from cluster 2 between the Garonne and the high-speed railway (Figure 5.C).

Although the development of the industrial area and the building of the high-speed railway do not directly threaten the *E. calamita* survival in the study site (all *p*_*e*_ = 0) (figure 6.F), it will restrain the species’ displacement capabilities and reduce the global population size (respectively, average population size within the focal patches are 2907, 2635 and 2166 individuals). In addition, figure 8.F highlights the central role of patch 50 as a stepping-stone patch, which allows the maintenance of connectivity between the north (patches 29 and 32) and the south (patches 71 and 103). Such a map constitutes a powerful decision-making tool in the new ecological context of compensatory measures which will be taken in the context of large-scale landscape planning (Lanius, Kiss & Den Betsen 2013; Regnery, Couvet & Kerbiriou 2013).

## Model limitations

By its structure, MetaConnect allows the user to take into account most of the requirements necessary for a complete and flexible PVA and decision-making tool (i.e. metapopulation dynamics and genetics) (Grimm *et al.* 2004; Baguette *et al.* 2013; Pe’er *et al.* 2013).

Currently, its main limitation comes from the landscape representation. In MetaConnect the landscape is represented using the patch-matrix approach, in which a cell is a suitable habitat or not. This approach will not be fully unrealistic for many species (Clobert *et al.* 2001; Urban *et al.* 2009; Pe’er *et al.* 2011). Furthermore, reproduction in the suitable habitat is assumed to be panmictic which is usually not true because patch shape and structure isolate or aggregate individuals within a patch and individual behaviour (territoriality, mating system, cooperation,…) may aggregate or isolate individuals within a patch (Doebeli & Koella 1994; Snyder & Chesson 2003; Ylonen, Pech & Davis 2003; Nonacs & Kapheim 2007). Further development of MetaConnect toolboxes would allow tackling this limitation of intra-patch spatial structuring by splitting individual use of space into daily movements and dispersal events (Mueller & Fagan 2008; Roshier, Doerr & Doerr 2008; Pe’er *et al.* 2013).

A second limitation is the way genetic mutation is modelled. Currently, the mutation model is very simple, just assuming that a new allele can occur at a given constant rate and that no reverse mutations are possible. Further toolbox development would allow various methods for modelling mutations to be incorporated (Willi, Van Buskirk & Hoffmann 2006; Neher 2013; Wray 2013) and in addition would permit simulation of the action of the genotypes on the individual phenotype (Montalvo *et al.* 1997; Mouquet *et al.* 2012; *Moulherat et al.* submitted).

## Conclusion: model application and perspectives

European regulation requires spatial planners to evaluate precisely the impacts of developments on ecological network functioning. Baguette et al. (2013) recommended a robust workflow in that direction. The procedure comprises performing an analysis of metapopulation dynamics and dispersal over a landscape for each affected species to design sound ecological network functioning. MetaConnect provides an important tool that moves in this direction. Indeed, the user can easily build consensus networks for several species within the same study site and under the same framework with standardized and comparable outputs. However, such an approach does not yet incorporate the inter-specific interactions that could greatly change population dynamics and dispersal (Caswell 2001; Clobert *et al.* 2013). Further MetaConnect toolbox development will integrate basic inter-specific interactions such as competition, predation and parasitism.

Application of graph-theoretic connectivity is increasing at an exponential rate in ecology and conservation (Kindlmann & Burel 2008; Kadoya 2009; Moilanen 2011). In this framework, graph nodes represent habitat patches and graph edges represent the connectivity between patches (Urban & Keitt 2001). Whilst the mathematical background of graph theory is promising to deal efficiently with the ecological connectivity (Urban & Keitt 2001), the simplification made by modelling a metapopulation with a graph (Moilanen 2011) has the potential to lead to limited interpretation and operational efficiency (Kadoya 2009; Moilanen 2011). Kadoya (2009) concluded that graph modelling of metapopulations provides little congruence with connectivity inferred from population genetic structure and Moilanen (2011) listed the limits of graph-theoretic connectivity in spatial ecology. With MetaConnect, as shown in our case study (figure 6.A-C), we provide an efficient tool to define habitat patches (nodes of a graph) with predictions of the demographic module and dispersal functionality between patches (graph’s edges) with dispersal assessed from dispersal simulation or derived from genetic estimates applicable from local to national scales and grid-based data. However, further development of MetaConnect and the development of a specific toolbox will allow that the graph does not model the metapopulation functioning as such, but to be used as a powerful analytical tool preventing the graph-theoretic connectivity analysis from biases described by Moilanen (2011).

We conclude by highlighting the recent call for a new generation of models that begin to provide predictive systems ecology (Evans, Norris & Benton 2012; Evans *et al.* 2013). This call argued that, while in a few sub-disciplines such as dynamic vegetation modelling and climate change modelling we have already developed a capacity for simulating complex systems, typically we lack such a predictive modelling capability elsewhere in ecological and evolutionary disciplines (Moorcroft, Hurtt & Pacala 2001; Evans, Norris & Benton 2012; Hoban, Bertorelle & Gaggiotti 2012; Purves *et al.* 2013; Bocedi *et al.* 2014). The climate modelling community and the dynamic vegetation modelling communities both possess several models with which they can conduct inter-model comparisons, conduct cross-validations, etc (Evans, Norris & Benton 2012; Pe’er *et al.* 2013). This is also now true of the population genetics community, which has several software packages available (reviewed in Hoban, Bertorelle & Gaggiotti 2012). We believe that MetaConnect, together with the recently published software RangeShifter (Bocedi *et al.* 2014) and the integrative statistical inferences procedure proposed by Pagel & Schurr (2012), begin to address this challenge to develop complex systems models that can be used to inform policy in the sphere of spatial and landscape ecology.

## Acknowledgement

SM, EB, JC and MB were supported by the European project SCALES funded by the European Commission as *Large-scale Integrating Project* within *FP7* under grant 226852. JC, JT, SP and MB were supported by EU FP6 Biodiversa ERANET TenLamas. JT and SP were supported by NERC. SM and MM were supported by the TerrOïko’s R&D internal program. This work is part of the “Laboratoire d’Excellence” TULIP (ANR-10-LABX-41).

## Where to find MetaConnect

MetaConnect (IDDN.FR.001.430001.000.S.C.2012.000.20600) is developed in C++ by SM and MM and runs exclusively in cloud-computing under UNIX. A Graphical User Interface (GUI) developed in OpenGL insures the access to MetaConnect from multiple platforms (currently available for Linux Centos7). MetaConnect is available for academic purposes contacting the authors (SM).

